# Rapid Sterilization of Clinical Apheresis Blood Products using Ultra-High Dose Rate Radiation

**DOI:** 10.1101/2024.12.14.628469

**Authors:** Stavros Melemenidis, Khoa D. Nguyen, Rosella Baraceros-Pineda, Cherie K. Barclay, Joanne Bautista, Hubert Lau, M. Ramish Ashraf, Rakesh Manjappa, Suparna Dutt, Luis Armando Soto, Nikita Katila, Brianna Lau, Vignesh Visvanathan, Amy S. Yu, Murat Surucu, Lawrie B. Skinner, Edgar G. Engleman, Billy W. Loo, Tho D. Pham

## Abstract

**BACKGROUND AND OBJECTIVES:** Apheresis platelets products and plasma are essential for medical interventions, but both still have inherent risks associated with contamination and viral transmission. Platelet products are vulnerable to bacterial contamination due to storage conditions, while plasma requires extensive screening to minimize virus transmission risks. Here we investigate rapid irradiation to sterilizing doses for bacteria and viruses as an innovative pathogen reduction technology.

**MATERIALS AND METHODS:** We configured a clinical linear accelerator to deliver ultra-high dose rate (6 kGy/min) irradiation to platelet and plasma blood components. Platelet aliquots spiked with 10^5^ CFU of *E.coli* were irradiated with 0.1-20 kGy, followed by *E.coli* growth and platelet count assays. COVID Convalescent Plasma (CCP) aliquots were irradiated at a virus-sterilizing dose of 25 kGy and subsequently, RBD-specific antibody binding was assessed.

**RESULTS:** 1 kGy irradiation of bacteria-spiked platelets reduced *E.coli* growth by 2.7- log without significant change of platelet count, and 5 kGy or higher produced complete growth suppression. The estimated sterilization (6-log bacterial reduction) dose was 2.3 kGy, corresponding to 31% platelet count reduction. A 25 kGy virus sterilizing dose to CCP produced a 9.2% average drop of RBD-specific IgG binding.

**CONCLUSION:** This study shows proof-of-concept of a novel rapid blood sterilization technique using a clinical linear accelerator. Promising platelet counts and CCP antibody binding were maintained at bacteria and virus sterilizing doses, respectively. This represents a potential point-of-care blood product sterilization solution. If additional studies corroborate these findings, this may be a practical method for ensuring blood products safety.

**HIGHLIGHTS:**

- 1 kGy irradiation of bacteria-spiked platelets reduced *E.coli* growth by 2.7-log without significant change of platelet count, and 5 kGy or higher produced complete growth suppression.
- The estimated sterilization (6-log bacterial reduction) dose was 2.3 kGy, corresponding to 31% platelet count reduction.
- A 25 kGy virus sterilizing dose to CCP produced a 9.2% average drop of RBD- specific IgG binding.

## INTRODUCTION

In the crucial domain of transfusion medicine, whole blood and its derivative products like red blood cells, platelets, and plasma stand as cornerstones of life-saving interventions. However, there are also life-threatening associated risks with blood transfusions.

Rates of transfusion transmitted infections (TTIs) by known viruses such as HIV, HBV, and HCV have been reduced over the decades through a combination of infectious disease screening and donor history questionnaire [1–3]. However, despite these improvements, there remains a residual risk, which refers to the chance of transmitting a virus from donor blood that has passed screening undetected, typically because the donor is in the window period of infection when the virus is not yet detectable by tests [3–5].

The rate of bacterial contamination due to the storage conditions of platelets at room temperature—a conducive environment for bacterial proliferation within the product’s shelf life can lead to life-threatening reactions; 1:10,000 for bacterially contaminated products, and 1:50,000 for septic transfusion reactions. Diversion pouches and better arm disinfection decreased the platelet contamination rate by up to 77% [6], while advanced methodologies like large volume delayed sampling (LVDS) for both aerobic and anaerobic culturing, pathogen reduction technology (PRT), as well as secondary methods such as Verax Biomedical rapid platelet PGD testing have further contributed to decreasing the infection rates. Particularly noteworthy within pathogen reduction technologies (PRT) are techniques relying on ultraviolet illumination in the presence of photoactive substances like amotosalen/psoralen [7] or riboflavin [8], which, despite their proven effectiveness, are not without potential risks of immune reactions or toxicity [9]. Although these advancements stand as testaments to the evolution of blood safety practices, the risks are yet not negligible.

In the US, Intercept is the only FDA-approved PRT method for platelets [10]. Although effective at reducing bacterial contamination, this method is laborious as it includes an adsorption step that removes extraneous amotosalen to prevent a reaction in the eventual patient recipient. Furthermore, with many blood centers having a dual inventory of LVDS and Intercept-PRT, the operational complexity becomes a pain point in platelet production. Therefore, developing a PRT method with a simple workflow and without the need of a small photoactive reagent would be extremely advantageous.

Application of irradiation to biological tissues is a standard sterilization practice for tissue banks. International Organization for Standardization (ISO) and the American Association of Tissue Banks (AATB) provide guidelines for various tissue types routinely irradiation for clinical use. Bone, connective tissue, skin, and vascular grafts commonly receive between 15 to 35 kGy. Corneas and amniotic membranes, are treated with between 10 to 25 kGy. These irradiation procedures are critical for reducing the risk of disease transmission, ensuring that patients receive safe and sterile medical tissues.

However, the only example of radiation being employed in transfusion is for transfusion-associated graft versus host disease (TA-GVHD), which is more common in neonatal intrauterine transfusion recipients, certain immunocompromised individuals, recipients of transplants from relatives or HLA-matched donors, and those who have had marrow or stem cell transplants. In this clinical scenario, the cellular target is the donor’s residual white blood cells in the donated product, and the irradiation dose is 25 Gy targeted at the center of the blood product using Food and Drug Administration (FDA)-approved X-ray irradiation cabinets. The radiation dose is being delivered at standard clinical dose rates of ∼ 0.1 Gy/s (Quastar™ x-ray emitter), which are low dose rates and making it infeasible to use to reach the antibacterial dose range.

In contrast, linear accelerators such as those used for cancer therapy can be configured to deliver radiation at dose rates several orders of magnitude higher than conventional x-ray irradiation cabinets. In this study, we explore the potential of ultra-high dose rate (UHDR) irradiation to deliver bacterial and viral sterilization dose range in blood samples in a matter of minutes. Preclinical studies, both *in vivo* and *in vitro*, have utilized UHDR irradiation due to its association with lower toxicity to healthy tissues or cell culture when compared to conventional dose rate (CONV) irradiation. In our study, we utilize UHDR due to speed at which kGy doses can be delivered. Our hypothesis posits that UHDR irradiation can rapidly sterilize bacteria without adversely affecting the production of clinical-grade apheresis platelet products. We also provide proof-of-concept that linear accelerator-based irradiation can be a practical point-of-care solution for sterilization of blood products.

We have configured a clinical linear accelerator to uniformly administer doses of radiation to multiple 2 mL blood samples at a rate of 100 Gy/s using a 16 MeV electron beam. Both apheresis-derived platelet products and COVID-19 convalescent plasma (CCP) were exposed to a graduated dosing regimen up to a maximum of 25 kGy. Our findings indicate that a dose of 1 kGy reduced *E.coli* growth by 2.7-log without significant change of platelet count, while the estimated sterilization (6-log bacterial reduction) dose was 2.3 kGy, corresponding to 31% platelet count reduction.

Additionally, administering a dose of 25 kGy to CCP samples over approximately 4.2 minutes showed a negligible impact on the structural integrity of proteins, as determined by the sustained ability of antibodies to bind effectively. Our proof-of-concept study demonstrates the capability of UHDR technology as a promising tool for enhancing the security of platelet transfusions and convalescence plasma across different medical scenarios.

## MATERIALS AND METHODS

### Irradiation

All blood products were irradiated with a clinical linear accelerator (Varian Trilogy) configured for UHDR irradiations, as previously described [11, 12]. For the irradiation of multiple samples of blood products at the same dose, a 14-slot holder of 2 mL cryovials (internally threaded, Corning™) was designed in Fusion 360^®^ (Autodesk, San Rafael, CA, USA). 3D computer-aided design (CAD) files were edited with Ultimaker Cura v.4.3.1 and printed with Ultimaker S5^®^ (New York, NY, USA) using polylactic acid (PLA). The sample holder is designed to allow the vials with the samples to be immersed under liquid and consequently allow to control the temperature of samples during irradiation (**Figure 1A-C**). The cryovials were irradiated at source-to-surface distance (SSD; surface of the holder) of 44.1 cm (**Figure 1D**) with 20 × 20 cm^2^ jaw opening (actual field size at 100 cm SSD), resulting in approximately 10 × 10 cm^2^ irradiation field. The beam profiles and dose homogeneity were evaluated using radiochromic film at 44.8 cm SSD (central depth of the tube) with 1 cm solid water build up (**Figure 1E, F**). Electron beam irradiation geometry parameters are outlined in **Table 1**. Bacteria spiked samples were irradiated at room temperature, however, samples above 1 kGy were irradiated with sample containers immersed in pre-chilled water to avoid rise of temperature from radiation. CCP samples were irradiated in a frozen state, immersed in dry-ice-cold isopentane (2’-Methybutane, Sigma-Aldrich, US), containing pieces of dry ice to maintain a consistent temperature of approximately -78.5°C throughout the duration of irradiation. At the conclusion of irradiation, samples were returned to dry ice, before being returned to -80 °C storage, prior to further processing.

**FIGURE 1.**
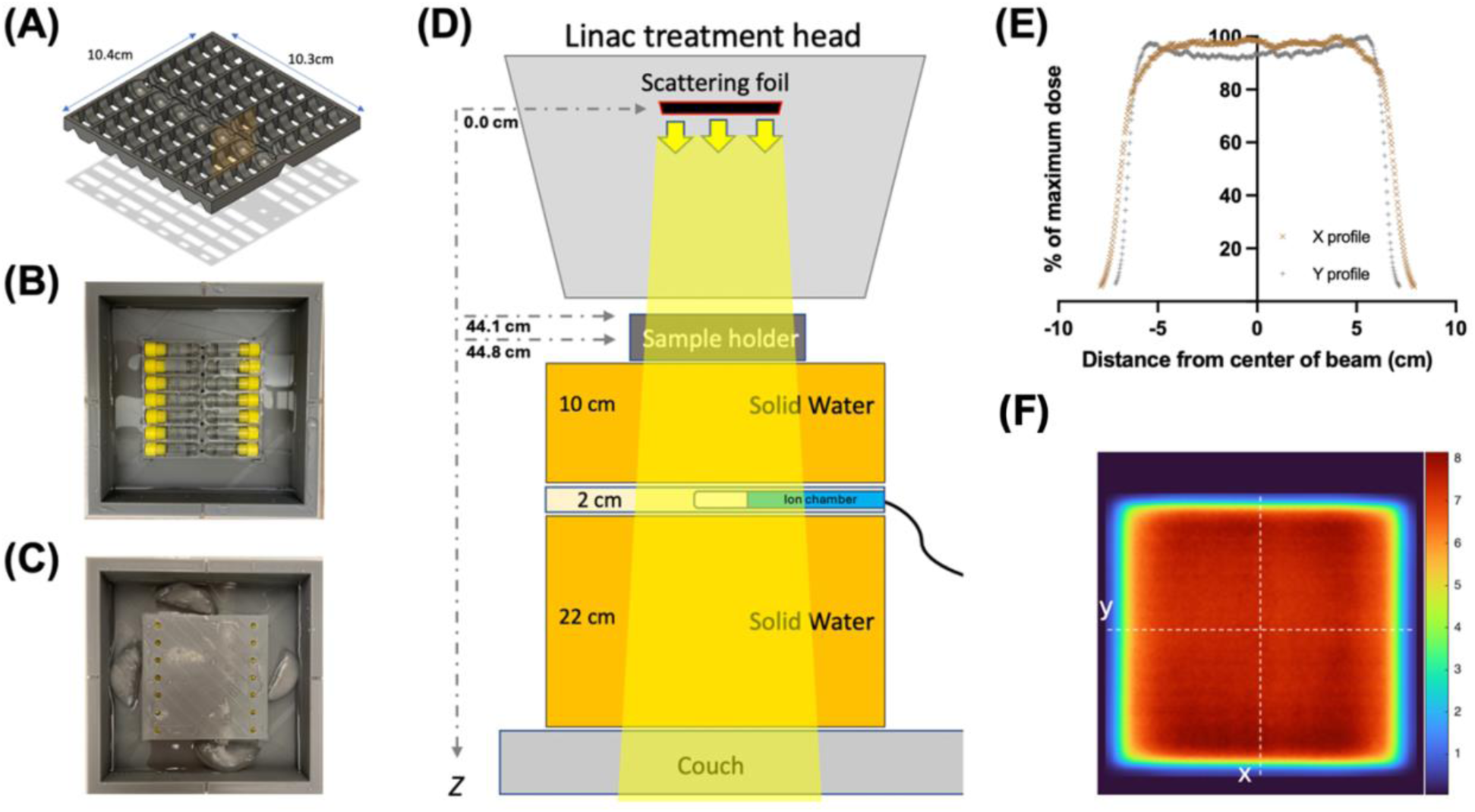
(**A-C**) Temperature controlled 14-slot blood sample holder for FLASH irradiation and (**D-F**) beam geometry. (**A**) CAD design illustrates a 14-slot holder for 2mL cryovial, that allows samples to be immersed in liquid (**B**), which can be used to control temperature. (**C**) Demonstrates samples at ∼4 *°*C. (**D**) Schematic diagram illustrates the geometry of the irradiation, with samples being at 44.1 cm source to surface distance (SSD; surface of the holder from scattering foil). The beam is monitored using an ion chamber (Farmer chamber) that measures the Bremsstrahlung tail of the electron beam. (**E**) The profiles of the beam were derived using film irradiated with 1 cm build up at 44.8 cm from the source (central depth of the tube) and are relatively flat and comparable in both x and y directions. (**F**) Dose map of the same films demonstrate relative homogeneity across the 10 *×* 10 cm^2^ area or interest.

**TABLE 1.** Electron Beam Irradiation Geometry Values and Experimental Parameters.

| Parameter | Value |
| --- | --- |
| Beam energy [MeV] | 16.6 |
| Source to surface distance (cm) | 44.8 |
| Field size (cm<sup>2</sup>) | 10×10 |
| Target dose [kGy] | 0.1, 0.5, 1. 5, 10, 15, 20, 25 |
| Pulse rate [Hz] | 180 |
| Average dose per pulse [Gy] | 0.53 |
| Average dose rate [Gy/s] | 95.5 |
| Pulse length [s] | 3.75E-6 |
| Average delivery time per kGy [s] | 10.44 |
| Intra-pulse dose rate [Gy/s] | 1.41E+5 |

### Dosimetry

The beam was monitored during irradiation by measuring the Bremsstrahlung tail of the electron beam using an ion chamber (Farmer® chamber 30010, PTW) placed at 59 cm SSD, upstream of the sample holder and 13 cm solid water. The measurements of charge were corrected for temperature and pressure. In order to calibrate the dose to blood at the depth of the center of the tube oriented perpendicular to the beam (**Figure 1B**), 2.4 × 5.1 cm^2^ pieces of radiochromic film (Gafchromic EBT-XD, Ashland™, Wayne, NJ, USA) were placed inside an empty holder filled with water. Films were analyzed as described previously [13]. A fixed number of pulses was delivered per film and, consequently, charges from ion chamber measurements were correlated to the associated absorbed dose from each radiochromic film, providing an average measured charge per dose (nC/Gy). The machine output was calibrated on a daily basis prior to each experiment. Electron beam irradiation experimental parameters are outlined in **Table 1**.

### Bacteria culture and preparation

Escherichia coli (ATCC 25922) obtained as a Culti-Loop culture (Thermo Scientific, Waltham, MA) was revived in 500 μL tryptone soy broth at 37°C for 5-10 minutes. The resulting bacterial suspension was streaked in the usual fashion onto blood agar plates and incubated at 37°C for 48 hours.

### Platelet components

Transfusable platelet component products were collected from volunteer blood donors (n = 5) at the Stanford Blood Center according to Association for the Advancement of Blood and Biotherapies (AABB) and FDA guidelines [14]. All platelet products were collected by automation. As part of the normal quality check (QC) process, after collection platelet products were transported to the lab where a platelet count was performed on each bag by Sysmex XN2100 instrument (Sysmex, Lincolnshire, Illinois). A small sample of these platelet products, with the platelet count, was aliquoted into 2 mL cryovials (Corning, Corning NY) that is to be sent for irradiation. One aliquot of 2mL was allocated for each irradiation group: 0 (non-irradiated control), 0.1, 0.5, 1, 5, 10, and 20 kGy. All aliquots were also spiked with approximately 10^5^ CFU of *E. coli* and stored at room temperature on a shaking incubator until irradiation. After irradiation, platelet counts were performed for each sample, and samples were immediately cultured on blood agar plates using a sterile calibrated loop at 37°C for 24 hours for colony counting.

### Plasma component

COVID Convalescent Plasma (CCP) products were also collected by automated method from volunteer blood donors at the Stanford Blood Center according to overall blood donor collection regulations, and additionally specifically according to FDA Guidance for the recruitment and collection of CCP [15]. At the time of collection, from each donor (n = 10) additional serum tubes were collected, aliquoted into 2 mL cryovials, and stored in -80 °C until needed for further experiments. For each of ten CCP donors, one aliquot was sent for irradiation in frozen state and the other was kept as frozen non-irradiated control. After irradiation, both sets were thawed and SARS- CoV2 antibody assays were performed to quantify any changes in binding activity.

### SARS-CoV2 Antibody-binding Assay

Previously described in detail [16], briefly here, we measured the antibody binding activity in the serum collected from healthy donors known to previously have been diagnosed with COVID-19. We employed an ELISA-based assay against Wuhan-Hu-1 SARS-CoV-2 (WT) Receptor Binding Domain (RBD) of the Spike protein as the target. 96-well plates were coated with 0.1µg per well of WT-RBD (ATUM) in PBS and incubated overnight at 4°C, then blocked with PBS-T containing 3% milk. CCP samples were diluted 1:1000, transferred to plates and incubated at 37°C for one hour.

Horseradish peroxidase conjugated goat anti-human IgG, IgA or IgM was used to detect isotype-specific binding. 100 µL of 3,3′,5,5′-Tetramethylbenzidine substrate solution was added to each well which developed for 12 minutes before 100 µL of 0.16 M sulfuric acid was added to stop the reaction. The optical density (OD) at 450 nanometers was measured with a SpectraMax M2 microplate reader (Molecular Devices, San Jose, CA).

### Statistics

For the dosimetry, the values presented are the averages of the measurements from all dose groups across multiple independent experiments (n=5). For the analysis of colony forming unit and platelet count assays, the dose groups (n = 5 donors per group) were compared to the non-irradiated controls using paired t-tests. A simple exponential decay fit was used to extrapolate the dose required for a 6-log bacterial killing. The percentage of remaining platelet count was estimated using a one-phase decay (exponential decay with plateau) fit. For the isotype-specific binding assays, each sample (n = 10 donors) was measured in duplicates. All values are presented as mean ± SD, except for the isotype-specific binding values, which are presented as the average of duplicates ± SD.

## RESULTS

### Irradiation and dosimetry

The profiles of the beam indicated reasonable flatness, with the sample holder receiving more than the 90% of the maximum dose in both x and y direction (**Figure 1E**).

Similarly, the dose map also confirmed relative homogeneity across the sample holder with constant dose above 90% of the maximum dose (**Figure 1F**). The average dose per pulse measured at the center of the tubes was 0.53 ± 0.02 Gy/pulse. The average time of delivery per 1 kGy was 10.44 ± 0.43 seconds, with an average of 1887 ± 77 pulses at 180 Hz pulse rate. The average dose rate was estimated to be 95.49 ± 3.69 Gy/s (**Table 1**).

### Irradiation dose-response on bacteria growth

Five platelet product aliquots per dose group from five healthy blood donor samples were spiked with approximately 10^5^ CFU/mL of *E. coli,* prior to irradiation. Samples from each donor were irradiated with a range of doses (0, 0.1, 0.5, 1, 5, 10 and 20 kGy), then plated for growth with CFU enumeration two days later. As shown in **Figure 2A** even 1 kGy was able to completely suppress bacterial growth for four out of five spiked samples, with one sample showing a 2-log reduction in bacterial growth. **Figure 2B** illustrates the average reduction of bacterial growth, which was a 2.7-log reduction from the initial seeding at 1 kGy. By extrapolation from an exponential decay fit, a 6-log bacterial killing dose considered to be sterilizing would be 2.3 ± 0.1 kGy.

**FIGURE 2.**
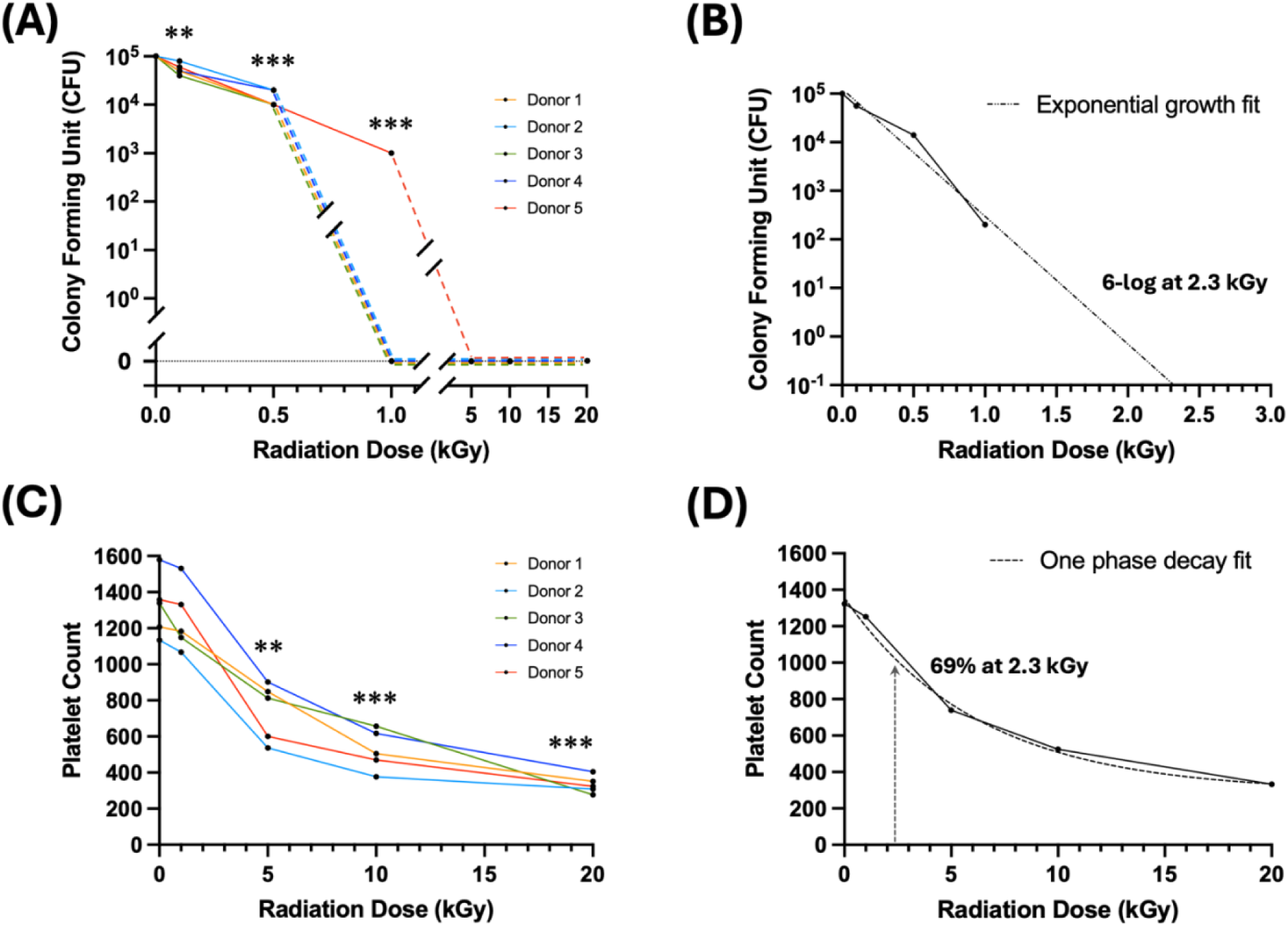
(**A,B**) Dose-response of *E.coli* bacterial survival and (**C-D**) platelet count from irradiated apheresis platelet products acquired from five healthy volunteer donors (one aliquot per dose per donor) spiked with approximately 10^5^ CFU of *E.coli*. (**A**) Bacterial survival in spiked platelet products post-irradiation per individual donor, and (**B**) averaged over all donors (dotted line is exponential decay fit). At 1 kGy dose only one sample (1 donor) showed 2-log bacterial reduction with no growth in the rest, while at 5, 10 and 20 kGy there was no bacterial growth in any sample. The extrapolated sterilizing 6-log killing dose is 2.3 ± 0.1 kGy. (**C**) Platelet counts in the same spiked platelet products post-irradiation per individual donor, and (**D**) averaged over all donors (dotted curve is one phase decay [exponential decay with plateau] fit). At 1 kGy dose, platelet count reduced minimally to 95% ± 5%. At estimated 6-log killing dose of 2.3 kGy the platelet count is estimated to be 69% ± 1%. Paired t-tests; *\*P* < 0.01; *\*\*P* < 0.005; *\*\*\*P* < 0.001; all comparisons are against non-irradiated control.

### Irradiation dose-response on platelet count

We also sought to determine the dose-response of platelet count over this dose range. We used the same spiked platelet product aliquots from the same donors, subjected to 1, 5, 10 and 20 kGy irradiation, including a non-irradiation control. Post-irradiation we evaluated the platelet count to assess the impact of the different irradiation doses. As shown in **Figure 2C** and **Table 2**, platelet count drops significantly from 1 to 5 kGy.

**TABLE 2.**
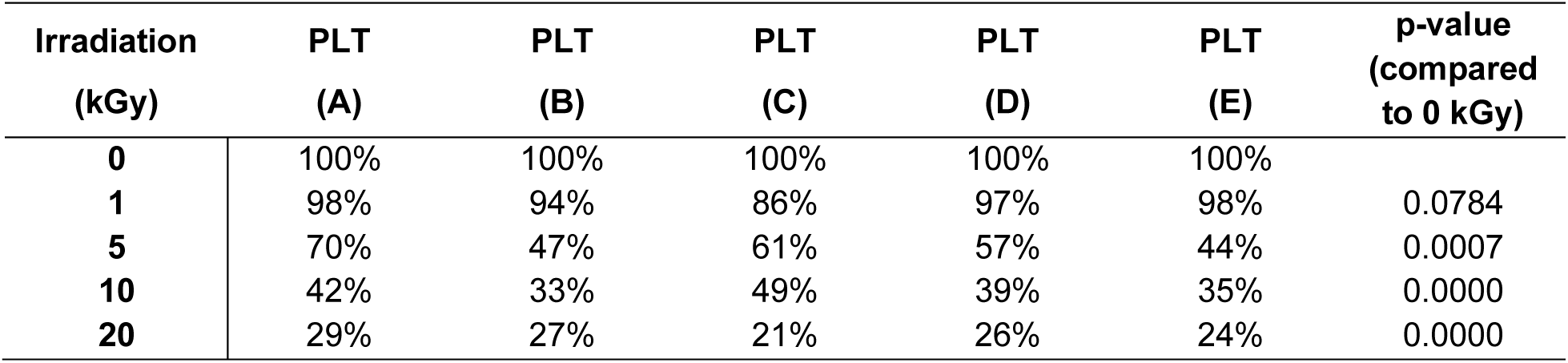
Dose-response of Platelet Count as Percent of Non-Irradiated Control.

Compared to the platelet count from unirradiated samples, 1 kGy radiation treatment reduced platelet count minimally to 95% ± 5% (*P* > 0.05), while 5 kGy reduced platelet count to 56% ± 11% (\*\**P* < 0.005) and 20 kGy to 25% ± 3% (*\*\*\*P <* 0.001). At the extrapolated 6-log bacterial killing dose of 2.3 ± 0.1 kGy for bacterial sterilization, the platelet count would be estimated to be at 69% ± 1% of the unirradiated controls (**Figure 2D**).

### Irradiation effects on protein integrity

Additionally, we sought to determine whether irradiation would have impact on protein binding structure in apheresis plasma products. Antibody binding in cognate antigen is highly dependent on an intact protein structure. Given that CCP is known to contain high levels of SARS-CoV-2-specific antibody, we investigated whether sterilizing doses of irradiation can abrogate the ability of IgG-mediated RBD binding in CCP. From ten CCP units of unique donors collected between March 2020 and June 2021, two aliquots of each were obtained for subsequent experimentation. One aliquot from each donor was used for 25 kGy irradiation, a dose chosen to be effective for viral sterilization; the second aliquot was kept frozen as the non-irradiated control. All samples were thawed and RBD-specific IgG antibody binding was determined by ELISA [16, 17]. As shown in **Figure 3**, RBD-specific IgG antibody binding showed a 9.2% drop on average. In contrast to IgG, changes to IgA and IgM mediated RBD-binding were not statistically significant (**Figure 3**).

**FIGURE 3.**
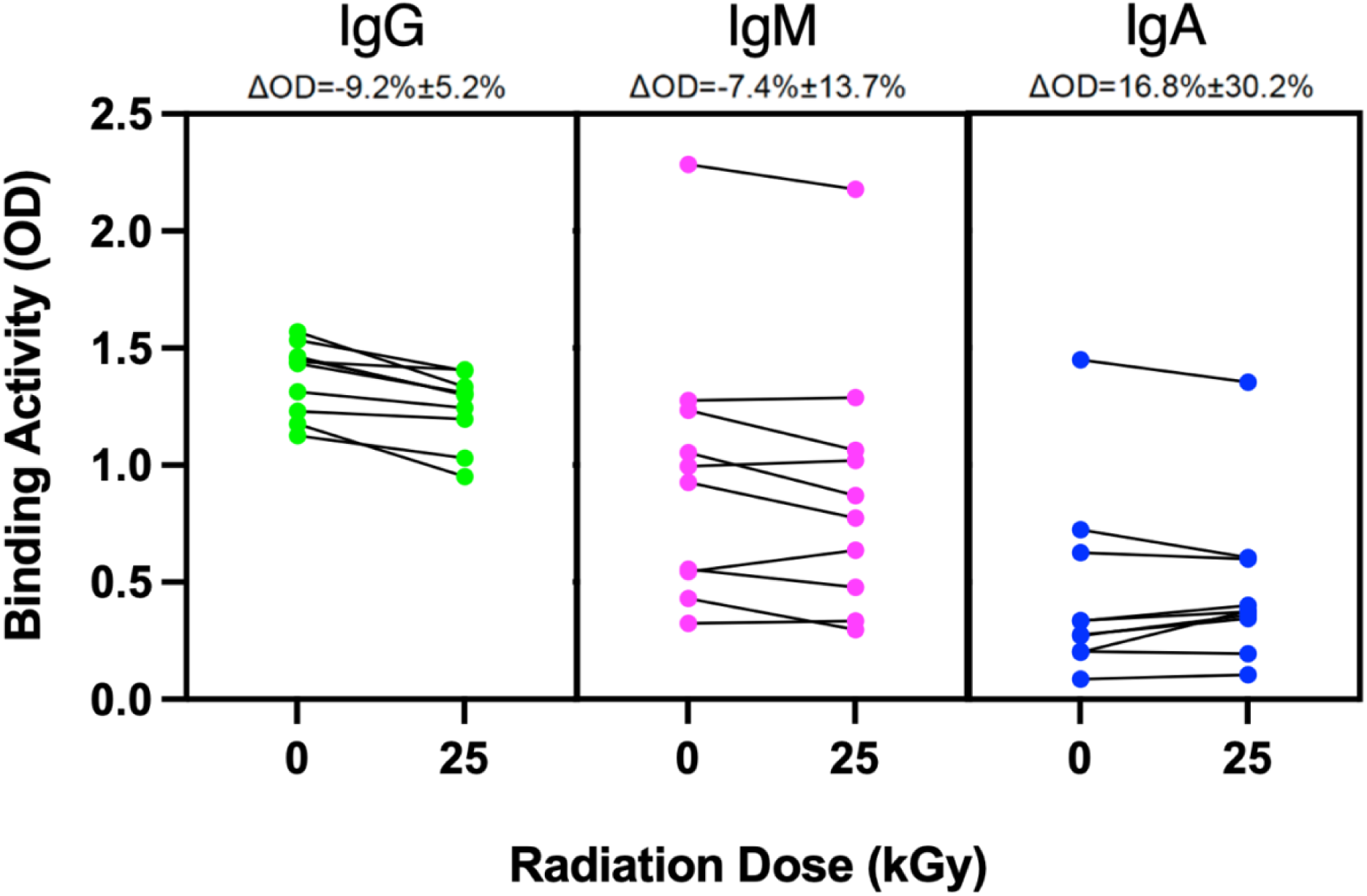
Effects of virus sterilizing dose of irradiation on SARS-CoV-2 RBD-specific antibodies in COVID Convalescent Plasma (CCP). 10 CCP donor samples were aliquoted for non-irradiated control and high-dose (25 kGy) irradiation. After irradiation, level of isotype-specific antibody binding was measured by ELISA, with OD405 readout. Each sample aliquot was done in replicate. Values represent average of duplicates ± SD for IgG, IgM, and IgA RBD-binding.

## DISCUSSION

We have described a novel method to reduce rapidly the bacterial load in apheresis platelet products collected from healthy volunteer blood donors. We could deliver 1 kGy irradiation in ∼10 seconds and 25 kGy in ∼3 minutes. In our pilot study, we have demonstrated that 1 kGy dose is high enough to reduce significantly bacterial growth from gram negative *E. coli* without significantly decreasing the platelet count in the product, and furthermore, that a 25 kGy viral sterilization dose only minimally decreased SARS-CoV-2 specific antibody binding.

Considering bacterial sterilization, we performed bacterial growth dose-response assays using *E. coli* as a pilot organism. Seeding at approximately 10^5^ CFU and with irradiation doses ranging from 0.1 to 20 kGy, we found that even 1 kGy induced complete bacterial growth suppression in 4 out of 5 spiked platelet samples at 48 hours. We estimate by exponential decay fitting that a sterilizing dose to produce 6-log killing is 2.3 kGy.

Conversely, we have determined the platelet count dose-response to irradiation in the range from 1 to 20 kGy. The platelet count did not significantly decrease for doses of 1 kGy or less. At the estimated 6-log bacterial killing dose of 2.3 kGy, the platelet count is reduced by only 31%, which is encouraging.

Moving forward, it is essential to replicate these experiments with a larger sample size and expand them to confirm more precisely that sterilizing doses for a range of pathogens can be administered while maintaining adequate platelet function.

To determine irradiation effects on protein integrity in plasma, we used CCP with known high titer of SARS-CoV-2 specific antibodies. Since an intact protein structure is crucial for antibody specificity, we assessed whether virus sterilizing doses of irradiation (25 kGy) would decrease RBD-specific antibody binding by ELISA. There was indeed a small but significant drop in IgG binding. However, we will need to determine whether this drop is considered significant in different clinical contexts. Although CCP had been a clinically relevant therapeutic in specific scenarios [18], now with vaccination and monoclonals widely available, as well as changes in COVID-related mortality, the small drop in RBD-specific RBD activity may be minimally impactful. However, to further determine effects of irradiation on protein integrity, it would be beneficial to measure changes in key proteins in the coagulation cascade related to increasing irradiation doses.

Further studies will also need to confirm sterilizing doses for clinically relevant transfusion-related viruses (*e.g.*, HIV, HCV, HBV, and WNV) as well as emerging infectious diseases for which there are currently no available tests (*e.g.*, dengue and chikungunya viruses). Because viruses generally possess smaller genomes relative to bacteria the probability of inducing a lethal lesion in each pathogenic organism per unit dose is lower, and the required dose to effectively sterilize viruses in blood products is substantially higher. Literature indicates that for 6-log viral inactivation 25 kGy appears to be on the upper end of the range for irradiation conditions above 0°C [19–25].

Moreover, since the mechanism by which irradiation damages cellular material is by radical formation, further exploration into the mitigating effects of antioxidants (*e.g.*, N-acetylcysteine) on blood product damage will be crucial, especially if the effects are different between blood components and pathogens.

The current work is presented as a proof-of-concept and requires additional future experiments to confirm and expand on these findings. Using irradiation for pathogen reduction is potentially simpler than current methods involving the addition, activation, and removal of small photo-active compounds. In fact, if eventually engineered appropriately, this usage would be analogous to current low-dose x-ray blood irradiation cabinets: using a small and enclosed chamber to irradiate the platelet product homogeneously before distribution or transfusion. Pathogen reduction can be done in matter of seconds for bacteria and few minutes for viruses without product manipulation, instead of hours with the current available PRT devices. Furthermore, it would be straightforward to design dedicated linear accelerator-based irradiators for practical implementation of this approach at the point of care in blood centers, in contrast to the large, centralized irradiation facilities used for industrial sterilization of medical products. We are hopeful that this new method of pathogen reduction may prove beneficial to the blood collection industry and transfusion services.

## AUTHORS’ CONTRIBUTION

All authors met the International Committee of Medical Journal Editors (ICMJE) criteria for authorship (https://www.icmje.org/recommendations/browse/roles-and-responsibilities/defining-the-role-of-authors-and-contributors.html). Stavros Melemenidis is the first author and author for editorial correspondence, and Tho D. Pham is the senior author and responsible for correspondence after publication.

## ACKNOWLEDGMENTS

SM, RM, MRA, EGE, MRA, MS, LBS, ASY, BWL, TDP: Conceptualization, Methodology; SM, RBP, CKB, JB, HL, VV, SD, NK, BL: Investigation, Data Curation; SM, BWL, TDP: Visualization; BWL, TDP: Supervision; SM, TDP: Writing – Original Draft; BWL, TDP: Writing – Review & Editing; SM, TDP: Project Administration; EGE, BWL: Funding Acquisition.

## FUNDING

This work was supported in part by the National Institutes of Health grants P01CA244091 (BWL) and R01CA233958 (EGE, BWL).

## CONFLICT OF INTEREST

BWL is a co-founder and board member of TibaRay. BWL is a consultant on a clinical trial steering committee for Beigene and has received lecture honoraria from Mevion. All other authors declare no conflicts of interest.

## DATA AVAILABILITY STATEMENT

The data sets used and/or analyzed during the current study are available from the corresponding author upon a reasonable request.

